# Straightforward semi-quantitative MALDI-TOF MS based screening approach for selection of recombinant protein producing E. Coli

**DOI:** 10.1101/2023.12.31.573754

**Authors:** Ivan N. Kravtsov, Andrey I. Solovyev, Eugenia A. Potemkina, Anastasya V. Kartashova, Maria A. Dmitrieva, Kseniya V. Danilova, Irina L. Tutykhina, Nikita B. Polyakov, Daria A. Egorova

## Abstract

MALDI-TOF MS represents a rapid and cost-effective method for identifying proteins and microorganisms. When obtaining recombinant protein producers, differences in expression levels among transformants necessitate the conduction of a small-scale screening of expression levels. This study proposes a fast and easy method for screening clones producing recombinant proteins using MALDI-TOF MS. Various recombinant proteins were utilized to test the proposed method

## INTRODUCTION

To obtain prokaryotic producers of recombinant proteins, small-scale experiments to test expression levels are recommended (Qiagen, June 2003). In addition to the influence of various cultivation factors, there is a scatter in the level of expression in individual transformants, which dictates the need for screening producers. Most formats for such testing involve growing individual clones in a small volume of medium, followed by lysis and verification using SDS-PAGE, Western blot, or dot blot. However, the use of instrumental methods for assessing expression levels is still extremely limited.

In this paper, we propose a quick and easy method for screening clones of recombinant protein producers using MALDI TOF MS. Our proposed approach, in addition to speed and scalability, also has significant potential for implementation in R&D.

In our study, we utilized sfGFP, bacterial proteins from the DNAIIb proteins group (The two subunits IHFa and IHFb of the Pseudomonas aeruginosa host integration factor (IHF) and the Staphilococcus aureus nucleoid-associated bacterial protein HU), their phage-derived inhibitor GP46 protein, and VHH (camelid heavy-chain-only antibody) antibodies as as recombinant protein models. The selected recombinant proteins for analysis differ in mass, their ability to form complexes with other bacterial cell molecules, and exhibit varying stability and toxicity to *E. coli*. For instance, in our research, obtaining IHFa and IHFb producers required extensive screening due to known issues with the stability of the IHFa subunit (Howard A. Nash, 1987).

Expression of the HU and GP46 proteins was not complicated by the properties of these proteins and did not require large-scale screening; however, the selection of clones using this protocol was carried out for reasons of speed and simplicity. The selection of VHH clones was previously carried out by classical methods, such as Western blotting; this work demonstrates the possibility of selecting clones with the maximum level of expression, which can be useful when using alternative technologies to phage display in which there is no selection of antibody variants adapted to bacterial expression. The sfGFP protein was chosen to demonstrate that it is possible to semiquantitatively assess the expression level of proteins up to 30 kDa without special sample preparation.

One of the main applications of MALDI-TOF mass spectrometers is a rapid and inexpensive approach for the identification of proteins and microorganisms. It is now available to many researchers and widely used in microbiology laboratories.

The availability of this device enables the optimization of research tasks and allows for the abandonment of some classical approaches.

## MATERIAL AND METHODS

### Plasmids and bacterial strains’

The genes for the bacterial proteins IHFa, IHFb, HU, the phage protein GP46, and the sfGFP gene were cloned into pET group plasmids. The gene encoding the VHH antibody was cloned into the pEXPR00 plasmid. To obtain producers of recombinant proteins, chemically competent cells of strain BL21 were transformed according to a standard protocol. (Sambrook, 2001)

**Table 1.**

| Protein | Vector | a.a. | Theoretical MW, kDa | Selected clone |
| --- | --- | --- | --- | --- |
| IHFa | pET | 109 | 12.536 | 45+45 |
| IHFb | pET | 105 | 11.881 | 50+50 |
| HU | pET | 103 | 10.980 | 25+25 |
| GP46 | pET | 87 | 10.153 | 10+10 |
| sfGFP | pET | 251 | 28.135 | 10 |
| VHH | pEXPR00 | 147 | 17.197 | 10+10 |

### Cultivation conditions

Luria-Bertani agar medium (Miller’s formulation) supplemented with appropriate selective antibiotics was used as the nutrient medium

The obtained transformants were spread onto Petri dishes and incubated at a temperature of 37°C for 12 hours.

Each isolated colony was subcultured at 37°C on a gridded master plate, both control and experimental. The nutrient medium of the experimental plate included Isopropyl-β-D-1-thiogalactopyranoside, with a final concentration of 0.5 mM in agar plates.

### Sample preparation

Four hours after incubating both control and experimental Petri dishes, individual daughter clones were collected from the surfaces of the agar in control and experimental dishes using a 10 µL plastic microbiological loop or a sterile pipette tip and transferred to the surface of a 386-spot steel target plate (Bruker Daltonics, Germany).

Then, 1 µL of 70% formic acid in water was added to each spot and dried at room temperature. The surface of the dried extract was covered with 1 µL of a matrix solution: a 20mg/ml solution of 2,5-dihydroxybenzoic acid (Sigma Aldrich, United States) in 30% acetonitrile with 0.5% trifluoroacetic acid (Panreac, United States), and it was dried at room temperature.

### MALDI-TOF MS analysis

Mass spectrometry analysis was performed on an UltrafleXtreme instrument (Bruker daltonics) equipped with a Nd:Yag laser (355 nm) in the linear mode. Positively charged ions were detected in the m/z range from 2000 to 30000 at the following settings of the ion source: voltage at IS1, 20 kV; at IS2, 19 kV; and amplification factor, 12.6. The spectra were recorded in automatic mode using the Flex Control program (v.3.4, build 135).

*Escherichia coli* DH5α with additional proteins (RNase A [M + H]+ 13683.2 Da, myoglobin[M + H]+ 16952.3 Da) protein extract served as a calibration standard (cat. no. 255343, Bruker Daltonics).

### Data processing

Spectra visualization was conducted using Flex Analysis 3.4 software package. After visual inspection, the data were transformed into a peak list for calculation and statistical analysis in GraphPad Prism 8.0.1

## RESULTS

### Specificity

The high sensitivity and precision of determining the molecular weight of proteins using the MALDI TOF method provide specificity, enabling the identification of recombinant proteins in producer strains (Fig. 2).

### Relative evaluation of recombinant protein production

The quantitative determination of protein concentration using the MALDI-TOF method is challenging due to variations in solubility, interaction with the matrix, and other physico-chemical characteristics. Consequently, there may be zones with different protein distributions within the matrix crystals in a single sample. To mitigate the impact of these factors, the spectra were recorded in the accumulation mode.

At the outset, to evaluate the capacity for assessing protein expression levels, transformants producing sfGFP were cultivated on a Petri dish with IPTG. These were then categorized into two groups of colonies based on their fluorescence intensity, which was assessed visually (Fig. 3).

After that, to evaluate the metrological characteristics of the linear model of the concentration and intensity ratios of peaks of target proteins, a number of dilutions of purified sfGFP protein dissolved in water (4, 2, 1, 0.25 ng/mkl) were directly applied to colony samples of the BL21(DE3) strain in three repetitions applied to the MALDI-TOPH target, then sample preparation was carried out according to the protocol provided in the materials and methods (Fig. 4).

The results obtained confirm the possibility of using this approach as a semi-quantitative method for assessing the level of expression and ANOVA results for linearity of the method

### High-scale screening and impact assessment

The use of expression constructs based on pET series vectors, regulated by the T7 phage transcription and translation system, involves IPTG induction. Following transformation with the respective vectors, clones were transferred to an agar master plate using a disposable microbiological loop, with and without an inducer. Following a 4-hour incubation, sample preparation and mass spectrometric analysis were carried out. In one set of experiments, peak intensities were normalized relative to RL33’s peak intensity (host cell) (Fig. 5) to assess the impact of heterogeneity in protein extraction from bacterial cells and their integration into the matrix.

The distribution of normalized and unnormalized peak intensity values ratios for target proteins in these samples is described by a power function (coefficient of determination 0.88) (Fig. 6). Thus, for semi-quantitative determination of expression levels, the normalization of values may be an unnecessary step and could result in an exponential increase in error as the intensity of the target peak increases.

## CONCLUSION

The characterization of expression profiles through the MALDI-TOF method is a widely accepted and well-established practice. Numerous approaches have been developed for the analysis of recombinant proteins, including the assessment of expression levels. However, certain multi-stage sample preparation techniques, such as centrifugation, biomass transfer, or methods that involve evaluating expression levels after chromatographic purification and enzymatic digestion of samples for quantitative assessment (Liu, 2022) may not be practical.

Compared to the standard MALDI Biotyper protocol (Elisabeth Nagy, 2012) and other protocols for mass spectrometric expression assessment (Michael L. Easterling, 1998), the proposed protocol (see Fig. 1) eliminates certain stages, such as growing clones in a liquid nutrient medium and sample preparation, which typically includes washing cells with water, lysis or extraction, and mixing with solvents and matrix in intermediate volumes (Serebriakova MV, 1999).

**Fig. 1.**
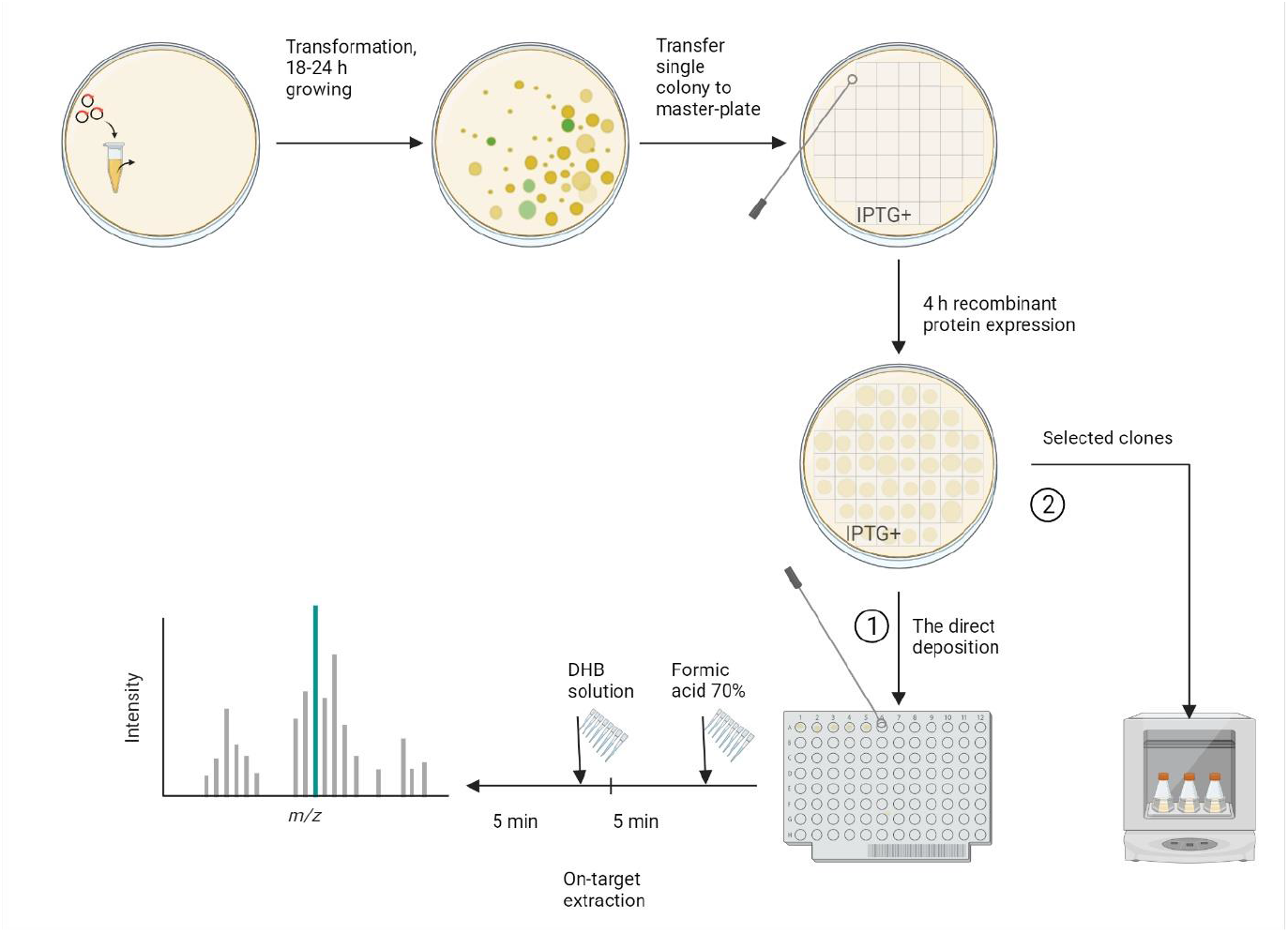
Workflows of the proposed clone selection protocol. (llustration was created in biorender). Each isolated colony was transferred to a gridded Petri dishes with agarose-containing LB supplemented with IPTG. After 4 hours, mass spectrometric analysis of the daughter clones was performed (1). Clones with the maximum intensity values of peaks of target proteins were then transferred from the master plate for cultivation in liquid nutrient medium (2).

**Fig. 2.**
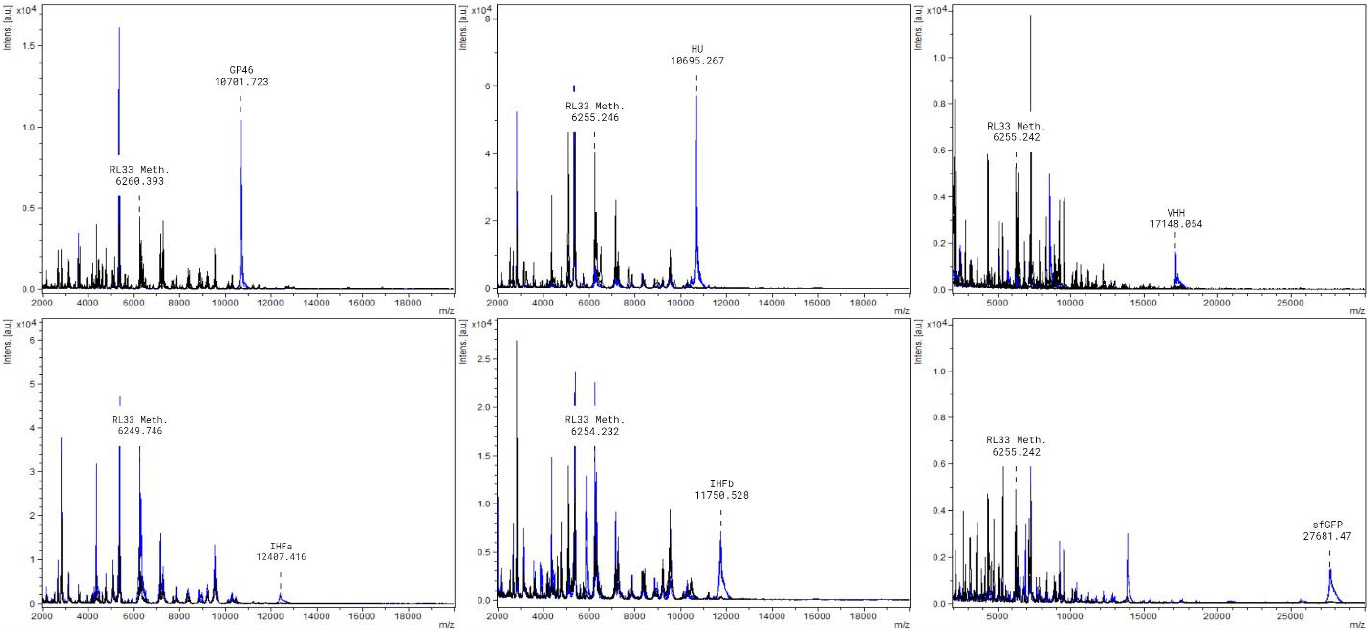
Comparison of mass spectra of colonies from producer strains and the control strain BL21(DE3). The spectra display the masses of peaks corresponding to recombinant proteins and the endogenous host cell protein -RL33(Meth.). Spectra from recombinant protein producers after IPTG induction are highlighted in blue, while control strains BL21(DE3) lacking an expression cassette are represented in black.

**Fig. 3.**
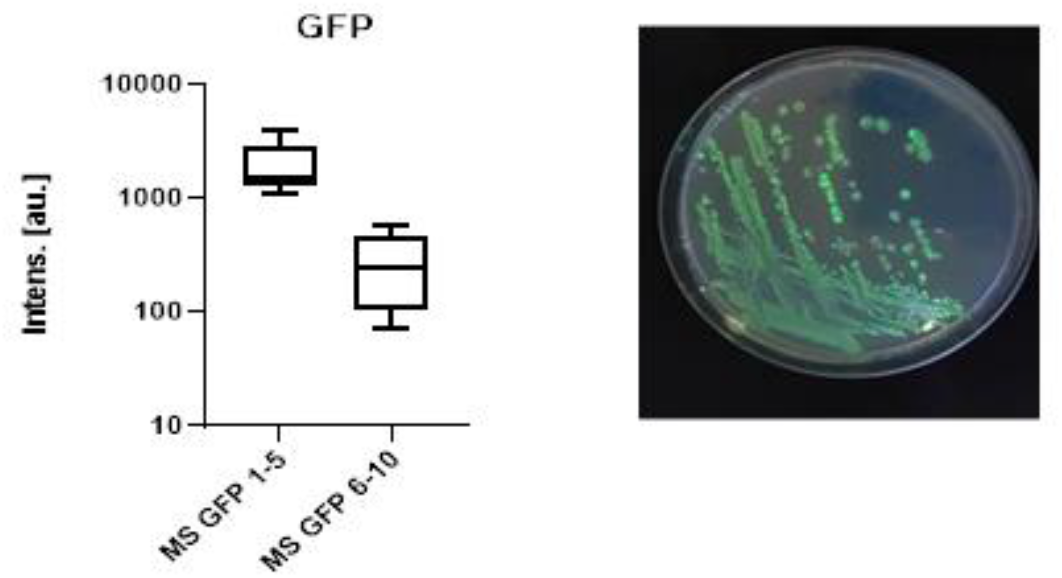
Intensity of the peak corresponding to the mass of sfGFP. MS GFP 1-5 is a group of colonies with high fluorescence intensity. MS GFP 6-10 is a group of colonies with low fluorescence intensity.

**Fig. 4.**
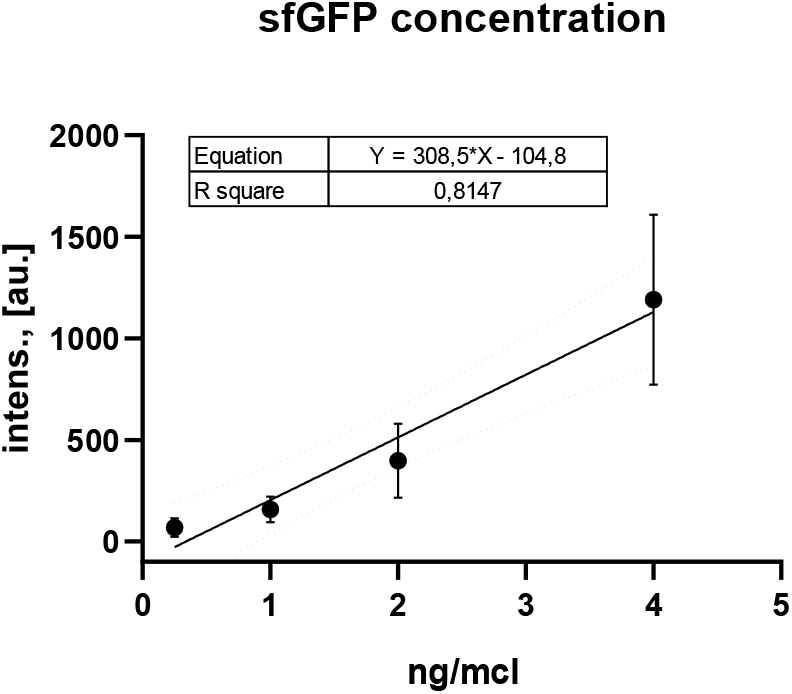
Ratio of concentrations to peak intensities of sfGFP

**Fig. 5.**
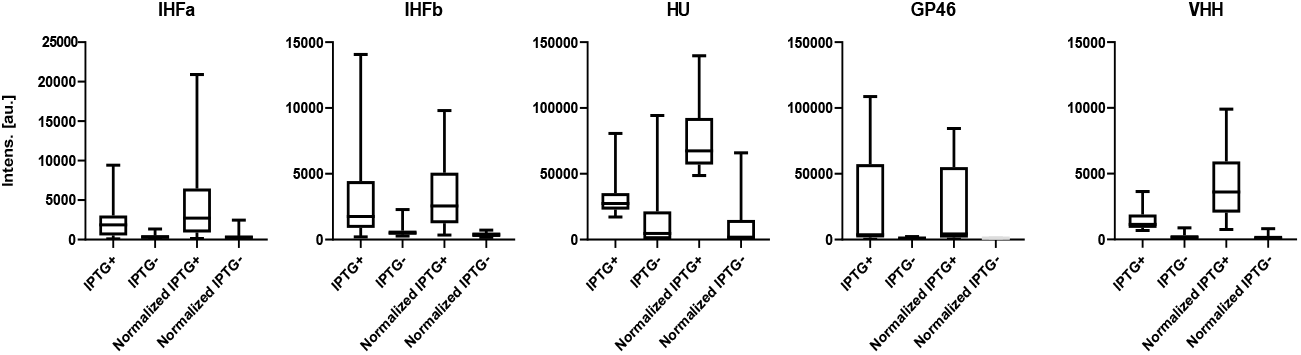
Distribution of peak intensities of induced clones and non-induced clones

**Fig. 6.**
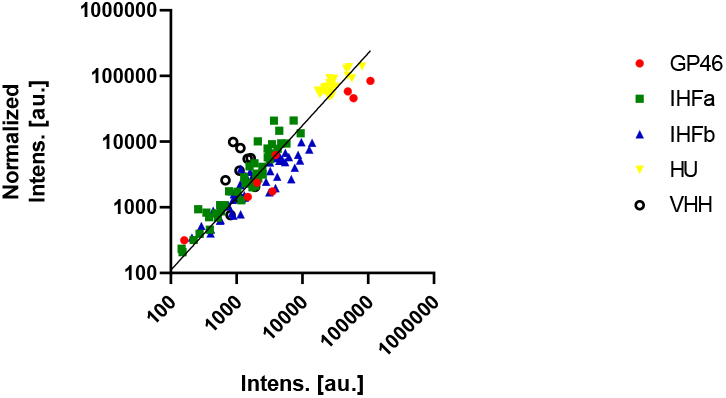
Ratio of normalized and non-normalized signal intensities

The use of Western blot and electrophoresis enables screening and semi-quantitative assessment of expression (Morão LG, 2022), but is constrained by the low throughput of the instruments and the availability of specific antibodies to the protein itself or tags, if applicable. Therefore, a rational approach is to employ Western blot to confirm the identity of the recombinant protein and the expression level in several clones after mass spectrometric screening.

The proposed method represents a compromise between the accuracy of expression level determination and the cost of analyzing a large number of samples. In our experience, the method allows for screening and selecting a producer from hundreds of colonies. Additionally, if necessary, such a system can be expanded and automated for high-throughput screening. In the future, we plan to test a mass spectrometric approach in high-throughput screening for the primary assessment of the productivity of eukaryotic producers, following a protocol with in-plate sample preparation.

In our study, this approach proved valuable for screening protein transformants that are toxic or have a significant impact on the physiology of E. Coli, such as IHFa. Additionally, the analysis of producers of recombinant single-domain VHH antibodies, widely employed in biomedicine, appears promising. Furthermore, fingerprinting the spectra of producer strains provides a easy and reproducible method for verifying species-specific characteristics and confirming the authenticity of qualities in deposited strains of genetically engineered producers used in the production of therapeutic drugs.

## Notes

### Competing Interest Statement

The authors have declared no competing interest.

